# An interactive deep learning-based approach reveals mitochondrial cristae topologies

**DOI:** 10.1101/2021.06.11.448083

**Authors:** Shogo Suga, Koki Nakamura, Bruno M. Humbel, Hiroki Kawai, Yusuke Hirabayashi

## Abstract

Outer and inner mitochondrial membranes are highly specialized structures with distinct functional properties. Reconstructing complex 3D ultrastructural features of mitochondrial membranes at the nanoscale requires analysis of large volumes of serial scanning electron tomography data. While deep-learning-based methods improved in sophistication recently, time-consuming human intervention processes remain major roadblocks for efficient and accurate analysis of organelle ultrastructure. In order to overcome this limitation, we developed a deep-learning image analysis platform called Python-based Human-In-the-LOop Workflows (PHILOW). Our implementation of an iterative segmentation algorithm and Three-Axis-Prediction method not only improved segmentation speed, but also provided unprecedented ultrastructural detail of whole mitochondria and cristae. Using PHILOW, we found that 42% of cristae surface exhibits tubular structures that are not recognizable in light microscopy and 2D electron microscopy. Furthermore, we unraveled a fundamental new regulatory function for the dynamin-related GTPase Optic Atrophy 1 (OPA1) in controlling the balance between lamellar versus tubular cristae subdomains.

## Introduction

Optimizing the metabolic capacity of mitochondria requires convolution of the inner mitochondrial membrane (IMM) to increase its surface area where substances are exchanged between the intermembrane space and the matrix (West et al., 1999). These highly convoluted IMM infoldings are called cristae. Decades of work has demonstrated that the entire set of five macromolecular complexes underlying the electron transport chain (ETC) and therefore mediate ATP production through oxidative phosphorylation (OxPhos) are concentrated in cristae (Davies et al., 2011; Friedman and Nunnari, 2014). Therefore, since variations in the shape of cristae determine the surface area of the IMM and diffusion of molecules, proper control of cristae structure is essential for cellular homeostasis. Transmission EM (TEM), especially electron tomography (ET), has played pivotal roles in describing the ultrastructural features of IMM organization because of its unparalleled resolution and revealed that cristae can be subdivided into lamellar and tubular subdomains (Jakobs et al., 2020; Pánek et al., 2020). However, due to the limited lateral and axial field of view, ET reports only part of cristae structures in each mitochondrion (Stoldt et al., 2018). Therefore, studying the molecular and physiological mechanisms regulating the structure-function relationship of cristae organization is still highly limited for technical and computational reasons.

The combination of serial sectioning and scanning EM (serial scanning EM; ssEM, SEM tomography) has been developed as an alternative approach for investigating fine structures in 3D. In particular, Focus Ion Beam-Scanning Electron Microscopy (FIB-SEM) lends itself well to the investigation of ultrastructural features in 3D because of its isotropic X-Y-Z resolution in the low nanometer range. Given that all membrane structures are stained in EM visualization, segmenting structures of interest from gray scale images is indispensable for segmentation and 3D reconstruction analysis. However, with the increased throughput of image acquisition in FIB-SEM and related 3D EM approaches comes an unsolved challenge with limited throughput in image segmentation.

Recent advances in the application of machine learning for image classification, object detection, and pixel segmentation in various types of images started to improve the efficiency of EM analyses. Even so, the wide range of mitochondrial shapes and complex 3D organization of their cristae structures have limited precise and automated segmentation of their ultrastructural features by thresholding-based methods even when assisted by the classical machine learning. However, among the machine learning approaches, deep-learning-based segmentation methods are transforming image analysis in biological studies because of their versatility and efficiency in segmenting a variety of cellular membrane structures (Moen et al., 2019). Particularly, variants of U-Net have been used successfully for segmentation of biological images (Falk et al., 2019). The segmentation of mitochondria using deep learning has improved recently (Liu et al., 2020; Xiao et al., 2018; Žerovnik Mekuč et al., 2020). While image calculation speed increased, laborious manual processing, such as generation of training data, proofreading of the analysis, and converting files for processing images across multiple software programs is still a significant limitation in EM analyses.

Herein, we developed a new integrated platform called PHILOW, equipped with the active learning for an efficient iterative training data generation, and a new 3D structure prediction algorithm using 2D training datasets. PHILOW drastically improves segmentation analysis and overcome previous limitations of 3D FIB-SEM segmentation and 3D reconstructions by (1) reducing the amount of human labor required for segmentation and (2) increasing the precision of segmentation allowing quantitative analysis detailed 3D structures of mitochondria and cristae in a cellular context. To demonstrate the potential of this approach, we performed a comprehensive analysis of mitochondrial and cristae structures of 135 control and 324 OPA1 deficient mitochondria which revealed novel roles for OPA1 in the regulation of cristae structures.

## Results

### Development of a Python-based human-in-the-loop iterative segmentation platform for 3D EM datasets

The implementation of deep learning has greatly increased the throughput of serial section EM analysis. However, as computer power increases, steps involving manual operations, such as generation of ground truth data sets and correcting predictions from deep learning-based segmentations are becoming the main rate limiting factor for ultrastructural cellular analyses. Amongst current limitations is a difficulty in ground truth generation due to inefficient choice of training data sets. In most ground truth generation, random crops of images are chosen for a training dataset, which leads to redundant annotations of similar structures. To circumvent this and achieve highly accurate predictions based on a minimal amount of manual segmentations, we developed a deep learning scheme by employing human-in-the-loop (HITL) learning (**Figures 1A and S1**). HITL learning starts from a small manually segmented training dataset, instead of generating the whole training datasets at once. After the first round of learning, the trained algorithm displays structures it did not learn. Then, training data corresponding to the unlearned structure are added for the next cycle. Iterations of this HITL-learning generate a model covering more comprehensive structures of target objects. This aids in avoiding redundant generation of training data for similar structures. However, implementation of this iterative scheme was hindered because of the time and labor required for format conversions, export and import cycles of data across different applications, and accompanying file management. Thus, to circumvent these roadblocks, we developed an open-source analysis platform called **P**ython-based **h**uman-**i**n-the-**lo**op **w**orkflow (PHILOW) (**Figure 1B**). Since the existing tools for iterative segmentations are either based on classical machine learning techniques such as random forest (Berg et al., 2019) or designed for segmenting 2D images (Ouyang et al., 2019), we built PHILOW on top of napari, a multi-dimensional image viewer for python, equipped with a GUI and workflows suitable for handling 3D data. PHILOW is made up of several components: annotation assistance, visualization and data management, and model training and inference functions. For annotation assistance and data management functions, we implemented orthogonal views, clickable training data selection, and data exporter and importer functions. This allows selection of training data by button-click while observing the 3D data and creating annotations. Training can be started via GUI on GPU-enabled machines or via Google Colaboratory. Additionally, we are able to train the model without transforming images between multiple applications. The prediction results of the dataset can be immediately visualized via PHILOW. During the iteration processes, the training data are automatically separated into original images and corresponding annotations. These seamless transitions enable a smooth iteration cycle for structure prediction. If the accuracy of the prediction results is saturated by iterative cycles, a final manual correction is performed directly on PHILOW. In addition, an active learning (AL) approach has been implemented in PHILOW to assist the iteration process by displaying prediction results in regions with low confidence. With AL, less accurate slices can be selected as effective training data for the next learning cycle (**Figures 1A and 1C**).

**Figure 1.**
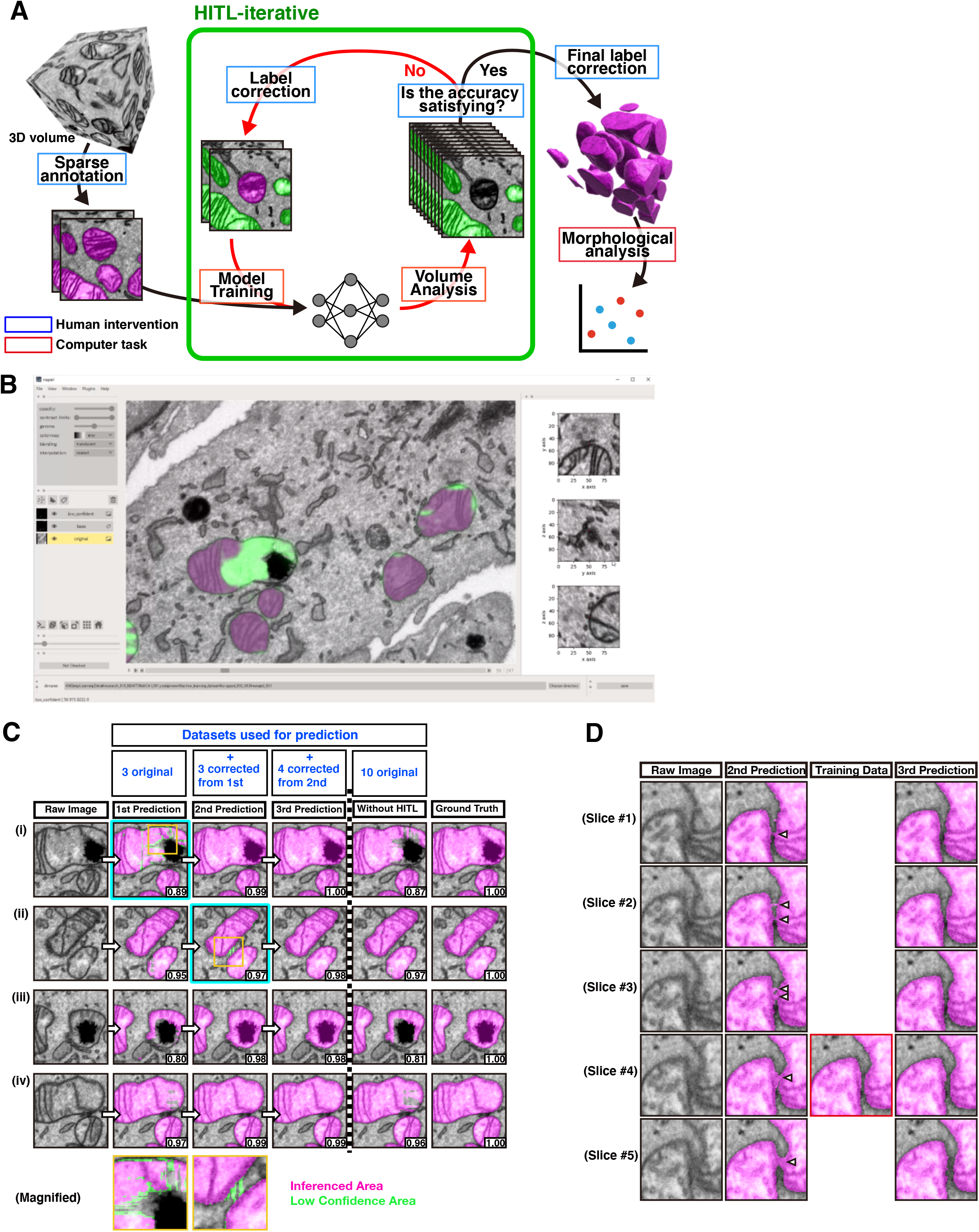
Implementation of HITL-iteration using PHILOW. **(A)** A diagram showing the human-in-the-loop iterative workflow. The green rectangle indicates processes performed iteratively with human intervention. **(B)** The graphical user interface of PHILOW during annotation. The buttons for data management and the orthogonal view are shown. Low confidence areas are highlighted in green for active learning. **(C)** Comparison of F1-scores between the HITL-mediated iterative learning and conventional deep-learning. Crops of raw image only (left), or overlaid with predictions for mitochondria (magenta) are shown. The F1-score of each crop is shown at the right bottom. In the HITL-mediated iterative learning, annotations of three randomly picked areas were used as an initial training dataset. After the first prediction, the annotations on three image crops, including image (i), were corrected and combined with the initial training dataset. F1-scores of second prediction with this new training dataset were improved not only in the image (i) but also in the image (ii) - (iv). Four image crops including image (ii) were corrected after the second predictions and combined with the training dataset for the second prediction. After these cycles, the F1-scores were above 0.98 in (i) - (v). In contrast, without HITL, even with the same number of training datasets, the accuracy reached only 0.81-0.97. Magnifications of the areas marked with orange rectangles are shown in the bottom. Low confidence areas were highlighted in green to draw attention of the annotators for the manual correction. **(D)** Segmentation results of mitochondria closely apposed to each other (magenta) are shown. The prediction mistakenly annotated those mitochondria as connected in the second prediction (arrowheads). At the third prediction, by adding a training data corrected from the second prediction in slide #4, misannotation in other slices were also corrected.

In summary, PHILOW allows us to display and correct prediction results, and apply the corrected results as training data for the next round of deep learning, all on a single platform.

### The human-in-the-loop approach enables efficient production of training data for deep learning and improves segmentation performance

To examine if the implementation of HITL increases the accuracy of predictions based on a given number of training data sets, we segmented mitochondria from FIB-SEM images of NIH3T3 mouse embryonic fibroblasts at 5×5×10 nm per voxel (**Video S1**). For a conventional prediction (without HITL), we generated a 2D UNet++ model by applying ten training image sets prepared from randomly selected areas (**Figure 1C**). For HITL-iterative prediction, we chose three random training data from the ten training data sets for generating an initial 2D UNet++ model. For the next cycle, we added three more training sets prepared by correcting inaccurate predictions from the first cycle (**Figure 1C**). For the third cycle, we added four more training sets based on the second prediction. Strikingly, while the quality of the predictions evaluated by the F1 score (the harmonic mean of precision and recall) were 0.909 on average by the model trained with the conventional method, with the constant increase of the F1 score along three iterative cycles, those by the HITL based model reached 0.97 (0.906 after the first cycle and 0.924 after the second cycle), even though the number of training data sets are the same (**Figure 1C**). The efficiency of HITL-based iterative method is underscored by the elimination of misconnected mitochondria, often seen in previously reported automated mitochondrial segmentations. Although the resolution of the images was sufficient for separating two closely apposed mitochondria, the predictions could not separate some mitochondria until the second iterative cycle (arrows in **Figure 1D**). Therefore, we included a training image that passed the correction of misconnections of mitochondria (Slice #4 in **Figure 1D**) in the next cycle. Strikingly, the third prediction separated the mitochondria precisely in serial sections. Thus, appropriate choice of training datasets by HITL significantly increases the performance of models without an increase in the volume of training data.

Next, to evaluate the effect of smooth HITL implementation using PHILOW, we measured total work time required for a complete 3D reconstruction of mitochondrial structures from 100 Mvoxel of 10 nm isotropic ssEM images (**Figure 2A**). Our target mitochondria appear in a variety of shapes from spherical to tubular as previously reported. Based on the time required for manual tracing of 0.9 Mvoxels, an estimated 3,546 minutes of human work would be required for a 100 Mvoxel manual reconstruction (**Figure 2B**). Using a 2D UNet++ model (DL baseline) trained with ten 640×640 pixel slices, the work time was reduced to 358 minutes (54 minutes for tracing of training data and 304 minutes of final correction). Strikingly, prediction by a 2D UNet++ model trained with three iterative cycles of HITL (HITL + DL baseline) reduced the work time for producing the training dataset to 38 minutes, while improving prediction accuracy. Therefore, manual correction took much less time (111 minutes). When we extended the analysis to neighboring image blocks, the efficiency was still better using HITL (1.18 minutes/Mvoxel, compared to 3.96 minutes/Mvoxel in DL baseline only), Therefore, as the database becomes larger, the total time required is 3 to 4 times shorter.

**Figure 2.**
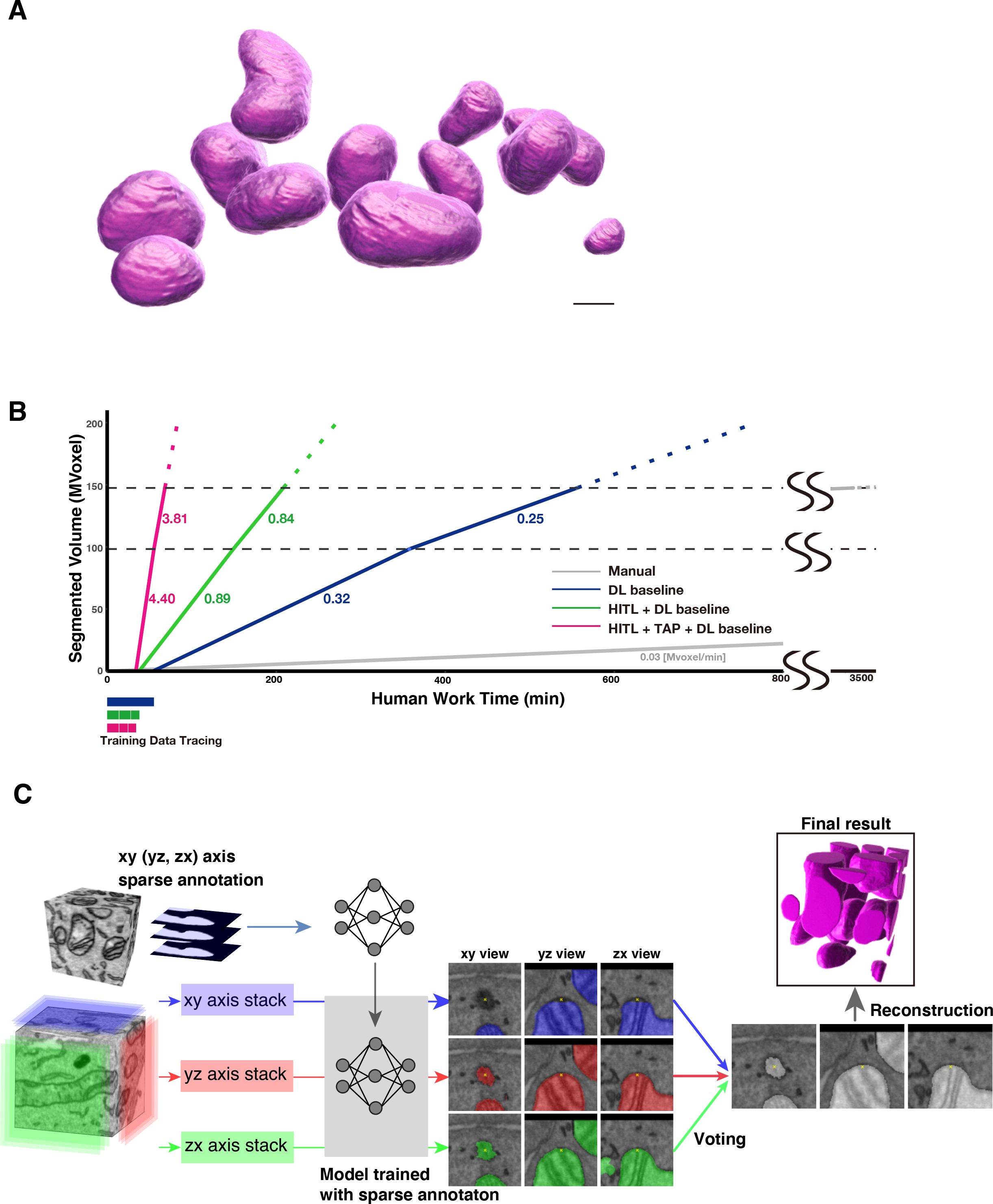
The HITL-TAP method on PHILOW improved the segmentation efficiency. **(A)** Representative 3D mitochondrial structures reconstructed from segmentations generated using HITL-TAP method on PHILOW. Scale bar, 500 nm. **(B)** Times required for correcting the mitochondrial prediction results obtained by indicated methods. Magenta: HITL + TAP + DL baseline learning. Green: HITL + DL baseline learning. Dark blue: DL baseline (2D-UNet++) only. Grey: without deep learning (Manual). The speed was calculated from the actual time required for correcting 150 Mvoxel (HITL + TAP + DL baseline, HITL+ DL baseline, and DL baseline) or the time estimated from 0.3 Mvoxel of manual correction (Manual). Bars below the graph show the time required for making the training data sets. Voxels visually inspected and corrected in one minute (Mvoxel/minute) either inside (0-100 Mvoxel) and outside (100-150 Mvoxel) of the volume used for generating the training data are indicated. **(C)** A diagram explaining the TAP method. A model trained with sparse annotation on xy plane images was applied for the virtual yz-(red) and zx (green)-plane images as well as xy (blue)-plane images. Therefore, each voxel had three prediction results from three axes. An example of three predictions visualized from three axes are shown in the middle. A majority vote was taken from the three predictions, and the result was adopted as the true value.

These results demonstrate that PHILOW-mediated seamless prediction and correction cycles significantly increased the efficiency of 3D reconstructions of organelle structures from ssEM images.

### The three-axis prediction based on 2D training data using PHILOW improves prediction accuracy of mitochondria structures

Although variations of 3D U-Net are applied for segmenting some types of 3D image volumes as a state-of-the-art algorithm (Lee et al., 2017; Xiao et al., 2018), the amount of human labor and experience required for generating a 3D training dataset is a major drawback. Therefore, we decided to create an algorithm to predict 3D structures by a deep learning model based on 2D training datasets. First, we predicted mitochondrial structures by a conventional 2D UNet++ model (Zhou et al., 2018). As expected, the predictions at the boundaries of mitochondria are frequently not accurate (**Figure S2A**). The section-to-section inconsistency at marginal areas increases variance of intersection over union (IoU) between segmentations of neighboring slices. To overcome this low accuracy, we developed a method called Three-axis prediction (TAP) that applies a 2D prediction model trained with a ground truth dataset from xy planes to virtual xz and yz planes (**Figure 2C**). First, we binned the 5×5 nm/pixel xy images to 10×10 nm/pixel for generating isotropic datasets with voxel size of 10×10×10 nm. Given that the voxel is isotropic, the model generated from the xy plane was directly applied to additional two planes and intermediate predictions from three axes were produced. Then, by majority vote of the intermediate predictions, the final prediction at each voxel was determined (**Figure 2C**). This enables a 3D inference without making extra training datasets. Strikingly, the IoU variance of predictions made by 2D UNet++ combined with TAP (TAP + DL baseline) was greatly reduced in xy planes (0.0082 compared to 0.0889 in 2D UNet++, **Figures S2A and S2B**). Further, because the prediction of mitochondrial structures using HITL + TAP + DL baseline is even more accurate than for HITL + DL baseline, the speed of manual correction (3.81 Mvoxel/minute) was more than 5 and 15 times faster than HITL+ DL baseline and DL baseline only, respectively (**Figure 2B**). This shows that the combination of HITL and TAP on DL baseline (HITL-TAP) enables highly efficient prediction of 3D structures from a total 10 sections of 2D training data that took only 33 minutes to prepare.

### Automated structural analysis of cristae using HITL-TAP

Studies using electron tomography (ET) have revealed that cristae consist of flat structures (lamellar cristae) and tubes (tubular cristae). Lamellar cristae are easy to observe because of their large and distinct structures, while tortuous and thin tubular cristae remain elusive. Given that 3D reconstruction using ET is limited to 200 nm thickness, reconstruction of cristae structures from serial FIB-SEM images is an attractive strategy to comprehensively observe cristae structure of each mitochondrion. Although several previous studies have manually reconstructed lamellar cristae structures (Pape et al., 2020; Stephan et al., 2020; Stoldt et al., 2018) manual reconstruction of the tubular structure has been challenging. To test if HITL-TAP on PHILOW can efficiently segment cristae structure including tubular structures from serial FIB-SEM images, we segmented the inner structures from 12 mitochondria that consist of a total 8,000 μm^3^ in 10 nm isotropic SSEM images (**Figure 3A**). 4 random images of 24 mitochondria were prepared as initial training data for both lamellar and tubular cristae to perform the initial prediction. Then, we carried out an iterative HITL cycle with TAP using three additional training datasets prepared by correcting the initial prediction. The total human work time was 33 minutes. The predicted data were compared with manual segmentations by two independent experts. For lamellar cristae, F1 scores between the prediction and segmentation of each human expert were comparable to that between the results of two human experts (**Figures 3B and 3C**). Close visual inspection revealed that the marginal areas of manual segmentations tend to include false positives as observed from yz- or zx- directions (**Figures 3D, 3E and S3**). This is represented by low precision scores of manual segmentations against HITL-TAP prediction (**Figure 3C**). Together with the smoother surface of prediction than the manual segmentation as shown by smaller IoU variances in xy-planes (**Figure 3F**), we conclude that lamellar cristae segmentation by HITL-TAP method has superior reliability compared to human experts. Of note, the prediction by the HITL-TAP method segmented a significant portion of tubular cristae that both human experts missed, as represented by the low recall scores (0.40 and 0.54) of human tracings against the prediction (**Figure 3C**). Post-hoc visual inspection in 3D (from yz and zx- planes) by the two experts confirmed that tubular predictions were indeed more efficient than manual segmentation (**Figures 3G-3J and S3**, arrow heads; **Video S2**). These results show that our HITL-TAP method achieved superhuman accuracy in segmenting both lamellar and tubular cristae structures (**Video S1**). This allows us to obtain previously inaccessible 3D cristae structures in whole mitochondrial network in situ.

**Figure 3.**
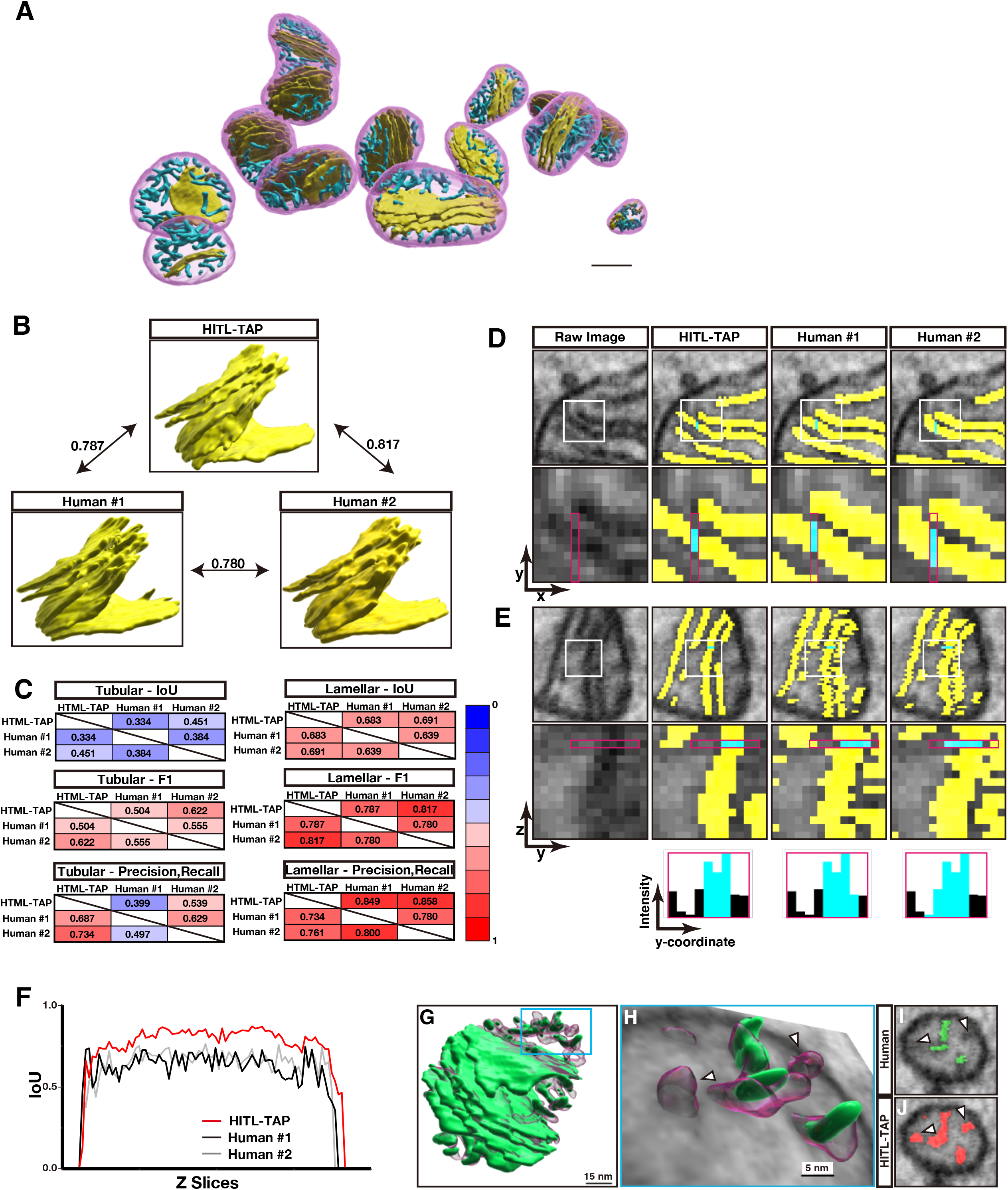
Prediction of cristae structures with superhuman accuracy. **(A)** Representative cristae structures in the mitochondria shown in Fig. 2a. Yellow: lamellar structure, Cyan: tubular structure. Scale bar, 500 nm. **(B)** Comparison of F1 score among a HITL-TAP prediction and annotations by two human experts. Corresponding 3D reconstructed lamellar images are shown. F1 score between the prediction by HITL-TAP algorithm and the annotation by human expert #1, and #2 are 0.787 and 0.817, respectively. This value is higher than the score between human experts (0.780). **(C)** Agreements among predictions by the HITL-TAP algorithm and annotations by two human experts were quantified by calculating IoU, F1 score, and precision/recall on tubular structures and lamellar structures. **(D and E)** Segmentation of lamellar structures by HITL-TAP algorithm and human annotators shown in xy planes (D) and yz planes (E). Rectangles in the upper panels show the areas shown in the lower panels. Yellow: segmentations of lamellar structures. Note that human annotators segmented false positive voxels at the edge of the lines highlighted in cyan. The continuity in the z-axis (e, original) implies the central three voxels are true positive. **(F)** Z-axis continuities of lamellar structures segmented by HITL-TAP algorithm (red), human annotator #1 (black) and human annotator #2 (grey) were indicated by IoU between neighboring xy planes. **(G)** Segmentation of cristae structures by HITL-TAP algorithm (transparent magenta) and a human annotator (green). The rectangle shows the area shown in (H). **(H)** Reconstruction of tubular cristae is shown with an EM slice. Note that the human annotation missed the structures HITL-TAP algorithm segmented (arrowheads). **(I and J)** yz-view EM images corresponding to the slice in (H) annotated by a human annotator (I) and HITL-TAP algorithm (J). Arrowheads are the spots corresponding to the tubular structures marked with arrowheads in (H).

### Quantitative analysis of 3D mitochondrial and cristae structures reveals that OPA1 is specifically required for tubular cristae structure

To demonstrate the potential of PHILOW-HITL-TAP methods for unbiased quantification of mitochondrial and cristae structures, we examined the roles of the dynamin-related GTPase Optic Atrophy 1 (OPA1). OPA1 was originally identified as a gene responsible for optic atrophy (Alexander et al., 2000; Delettre et al., 2000). OPA1 mediates IMM fusion and regulates cristae junctions (CJs), where cristae and the inner boundary membrane (IBM) are connected (**Figure 4A**; Barrera et al., 2016; Cogliati et al., 2016; Frezza et al., 2006; Harner et al., 2016; Olichon et al., 2003; Patten et al., 2014; Song et al., 2009). Given that previous studies assessed cristae structure in OPA1 deficient cells using either 2D EM images or ET, many of the structural phenotypes described, such as the continuity, orientation, volume, and surface area, were limited spatially. Further, the majority of analyses lack quantitative power.

**Figure 4.**
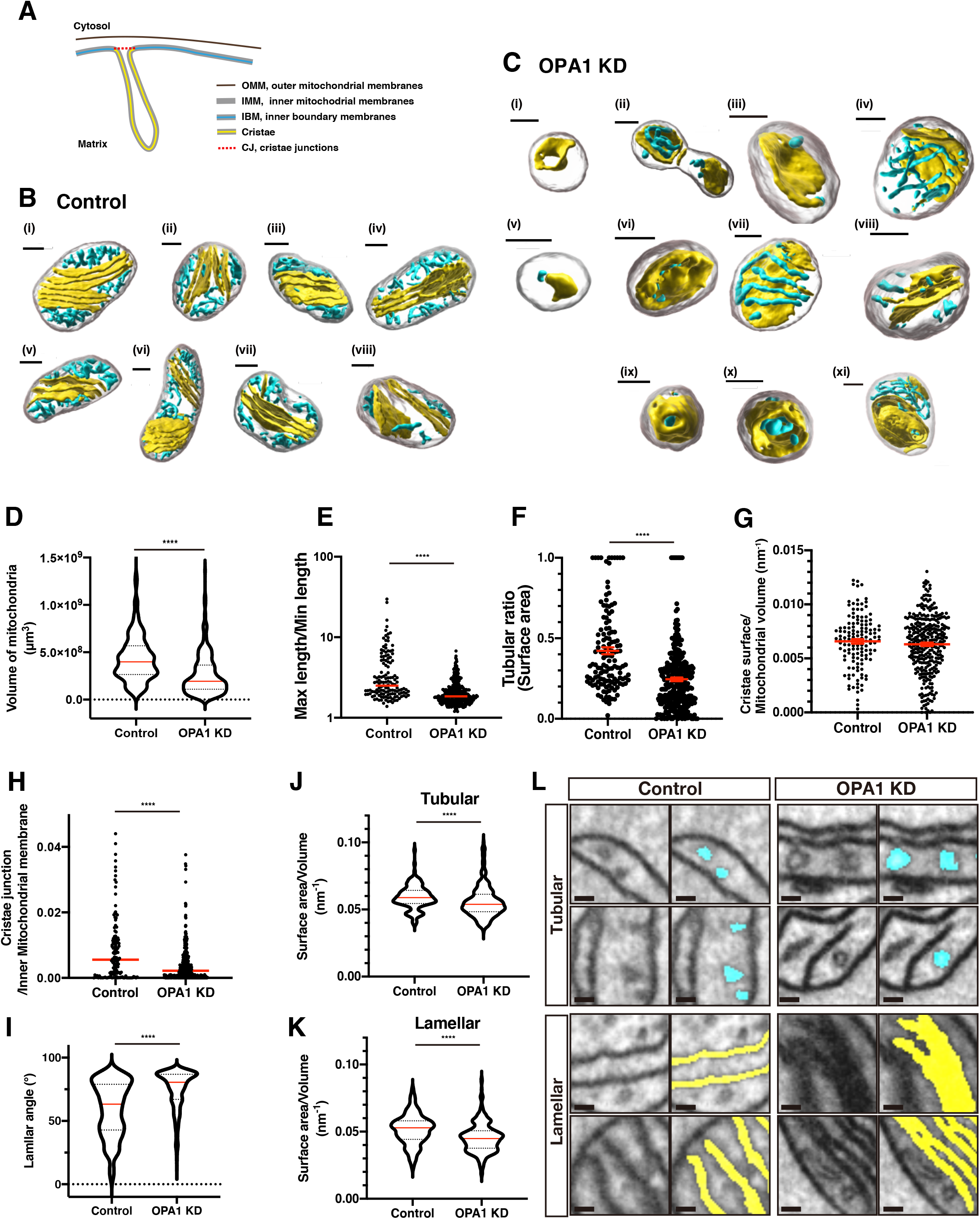
Quantitative analyses of the control and OPA1 KD cristae structures. **(A)** Diagram of mitochondrial subdomains. **(B and C)** Representative cristae structures of the control (B) or OPA1 KD (C) mitochondria. Scale bar; 300 nm. **(D and E)** Statistical analyses of the control and OPA1 KD mitochondria. Volume (D) and maximum length per minimum length (E) are shown with median (red lines). ****p < 0.0001, Mann–Whitney test. **(F-K)** Statistical analyses of the control and OPA1 KD cristae. Ratio of tubular cristae surface area to the total cristae surface area (F) and surface area of cristae per mitochondrial volume (G) are shown with mean ± SEM. The ratios of cristae junction area per inner membrane area in each mitochondrion are shown with median (H). One data point (0.073) is outside of the axis in the control. Angle between the vertical vector of lamellar cristae and the longitudinal vector of mitochondria (I). The average of the angles was calculated for each mitochondrion. Note that the angles are diverse in the control, while most of the angles are concentrated close to 90 degrees in OPA1 KD. Surface area per volume of tubular (J) or lamellar (K) cristae are shown with median. ****p < 0.0001, Mann–Whitney test. **(L)** Representative EM images of tubular and lamellar cristae from the control and OPA1 KD cells. Note that both lamellar and tubular cristae structures were thicker in the OPA1 KD mitochondria. Scale bar 100 nm.

To this end, NIH-3T3 cells, infected with lentivirus expressing either control shRNA or shRNA against OPA1, were imaged by FIB-SEM at 5×5×10 nm resolution. Then, xy-planes were binned to produce 10×10×10 nm isotropic images. Using HITL-TAP on PHILOW, we segmented outer and inner structures of 135 control and 324 OPA1 knocked down (KD) mitochondria from 5 cells each (**Figures 4B, 4C and S4A; Videos S3 and S4; Table S1**). Consistent with previous observations of fragmented mitochondria in OPA1 KD cells (Cipolat et al., 2004; Cogliati et al., 2013; Ishihara et al., 2006; Olichon et al., 2003), the median volume of OPA1 KD mitochondria was significantly reduced compared to control (**Figure 4D**, Control: 0.40 μm^3^, OPA1 KD: 0.19 μm^3^). The maximum length over minimum length ratio of the mitochondria in OPA1 KD cells were significantly smaller compared to the control, indicating that OPA1 deficient mitochondria were more spherical (**Figure 4E**).

Next, we analyzed the cristae structure of control cells. Previous observation of mitochondrial cristae in fibroblasts showed that the lamellar structure is dominant (Agier et al., 2012; Siegmund et al., 2018). However, our quantitative analysis showed that the surface area of the tubular structure as a percentage of the total surface area in each mitochondrion varied widely, spanning 10% to 100% with a mean of 42% (**Figure 4F)**. This suggests that the amount of tubular structure was vastly underestimated in previous studies. Importantly, we found that the ratio of tubular structure was significantly lower in the OPA1 KD mitochondria (mean 25%, p<0.0001, **Figure 4F**). This suggests that OPA1 is required for efficient formation of tubular structures at the expense of lamellar cristae topology. Interestingly, the total surface areas of cristae per 1 μm^3^ of mitochondria were not significantly different between the control and OPA1 KD cells (**Figure 4G**, Control: 6.6 μm^2^, OPA1 KD: 6.3 μm^2^; p=0.31). This indicates that OPA1 determines the ratio of tubular vs. lamellar structures while leaving the cristae area constant.

The regulation of CJ is important for cytochrome c release upon apoptosis and diffusion of ions and metabolites (Mannella et al., 2013; Scorrano et al., 2002). Although previous 2D EM studies have shown that the number of cristae with CJ was reduced in OPA1 deficient mouse adult fibroblast (Glytsou et al., 2016), whether the area and number of CJ per mitochondria is regulated by OPA1 was not investigated in these previous studies. Strikingly, the CJ area per IMM was reduced by 60% in OPA1 KD mitochondria **(Figures 4H and S4B)**. The number of CJ per IBM was also reduced (9.1/μm^2^ and 5.2/nm^2^ in the control and OPA1 KD, respectively; p=2.8×10^-6^). Indeed, large perforations of the lamellar structures are often observed in the OPA1 KD mitochondria (**Figures 4C-i, iii, v, vii)**. These results suggest that OPA1 is required for the proper formation of CJ in the mouse embryonic fibroblasts.

Next, we examined the orientation of cristae structures. Even though previous studies have observed that lamellar cristae are frequently aligned perpendicular to the longitudinal axis of the mitochondria (Wolf et al., 2019), this has never been analyzed in a quantitative manner. To examine if this is the case, we quantified the angle between the direction of the mitochondrial longitudinal axis (mitochondrial orientation) and the axis perpendicular to the lamellar cristae plane in 3D. Therefore, if the angle is zero, the lamellar cristae are perpendicular to the mitochondrial orientation. Although we observed lamellar cristae that clearly aligned perpendicular to mitochondrial orientation as observed with super resolution microscopy **(Figures 4B-i, iii, vi)**, a significant number of cristae were aligned parallel to the mitochondria orientation **(Figures 4B-iv, vii)**. Indeed, the average angle to the mitochondrial orientation was around 63° in the control (**Figures 4I and S4C**). This suggests that cristae orientation is near random in control mitochondria, contrary to the traditional view. On the other hand, OPA1 KD cristae had a marked tendency to be aligned in a direction parallel to the mitochondrial orientation (**Figures 4C-vi, viii and 4I**, average 81°). This is in line with a previous report describing the orientation of lamellar cristae in the OPA1 KD mitochondria (Griparic et al., 2004). This argues that OPA1 plays a role in connecting the mitochondrial shape and the formation of cristae junctions.

The surface per volume ratios of both lamellar and tubular cristae were significantly higher in OPA1 KD cells (**Figures 4J and 4K**). We hypothesized that this is due to the spherical morphology of OPA1 KD cristae. Indeed, in the OPA1 KD cells, multiple lamellar cristae were frequently segmented as connected structures, either because they were fused or the gaps between them were too close to be recognized (**Figure 4L**). Furthermore, the diameter of the cross section at the tubular cristae was clearly larger than the corresponding structures in the control (**Figure 4L**). These results are consistent with previous reports showing that cristae structures become thicker in the absence of OPA1 (Cogliati et al., 2013; Frezza et al., 2006).

It has been postulated that septa of inner membranes, which separate mitochondria in two compartments, formed in OPA1 KD mitochondria as a result of incomplete inner membrane fusion (Harner et al., 2016; Kojima et al., 2019; Sesaki et al., 2003). However, to our knowledge, a complete septum structure has never been shown. Using a septa-detecting algorithm (see figure legend for Figure S4D) and following visual inspection, we found 4 complete septa among 324 OPA1 KD mitochondria (**Figure S4D**). Since none of the 135 control mitochondria possessed a septum, OPA1 KD may induce septum formation.

Previous reports proposed that a section of an extended curved septum appears as an onion-like structure in the OPA1 KD mitochondria by stereological analysis (Harner et al., 2016). Indeed, we observed onion-like structures in 2D sections of OPA1 KD mitochondria (**Figure S4E**). However, reconstruction of those structures revealed that they are part of cristae mostly disconnected from the IBM and folding inside of the mitochondria (**Figures 4C-ix, xi; Video S5**). The isolation from IBM suggests that the onion-like structure represents a deficiency in the cristae junction formation rather than inner membrane fusion. This suggests that the onion-like structure is not a part of septum and rather resembles the cristae structures observed in *C. elegans* deficient for the OPA1 homolog eat-3 (Kanazawa et al., 2008). This finding emphasizes the advantage of our comprehensive 3D analysis of cellular ultrastructure to identify new cristae structure phenotypes and more generally improve our understanding of the molecular mechanisms regulating the structure/function relationship of mitochondrial physiology.

### Unsupervised learning of 3D cristae structures

To analyze mitochondrial ultrastructure systematically, given the large number of ultrastructural segmentation of mitochondria, we performed unsupervised learning of 3D morphological features of mitochondria and cristae. We used a six-parametric principal component analysis (PCA) to categorize mitochondrial ultrastructural features in an unbiased way (**Figures 5A and S5A-S5D)**. The parameters of mitochondria in control cells were used to fit the model, and the resulting model was subjected to dimensionality reduction algorithms for all mitochondria, including those in OPA1 KD cells. The volume and surface area of mitochondria and cristae contributed significantly to PC1, while the ratio of the tubular/lamellar cristae contributed to PC2. As a result, mitochondria of OPA1 KD cells showed a significantly distinct distribution compared to control (**Figure 5A**). This analysis showed that the ratios of tubular and lamellar cristae were not significantly different between the control and OPA1 KD mitochondria in a population within the range of −1≤PC2<1.5 (**Figures 5A-i and S5E**). On the other hand, another population of mitochondria with PC2 score higher than 1.5, is highly enriched in OPA1 KD cells that were smaller and had very fewer tubular cristae than control and lamellar cristae with irregular structures (**Figure 5A-ii**). These results suggest that PCA based classification based on ultrastructural features of mitochondria can distinguish OPA1 KD from control mitochondria (**Figure 5A**).

**Figure 5.**
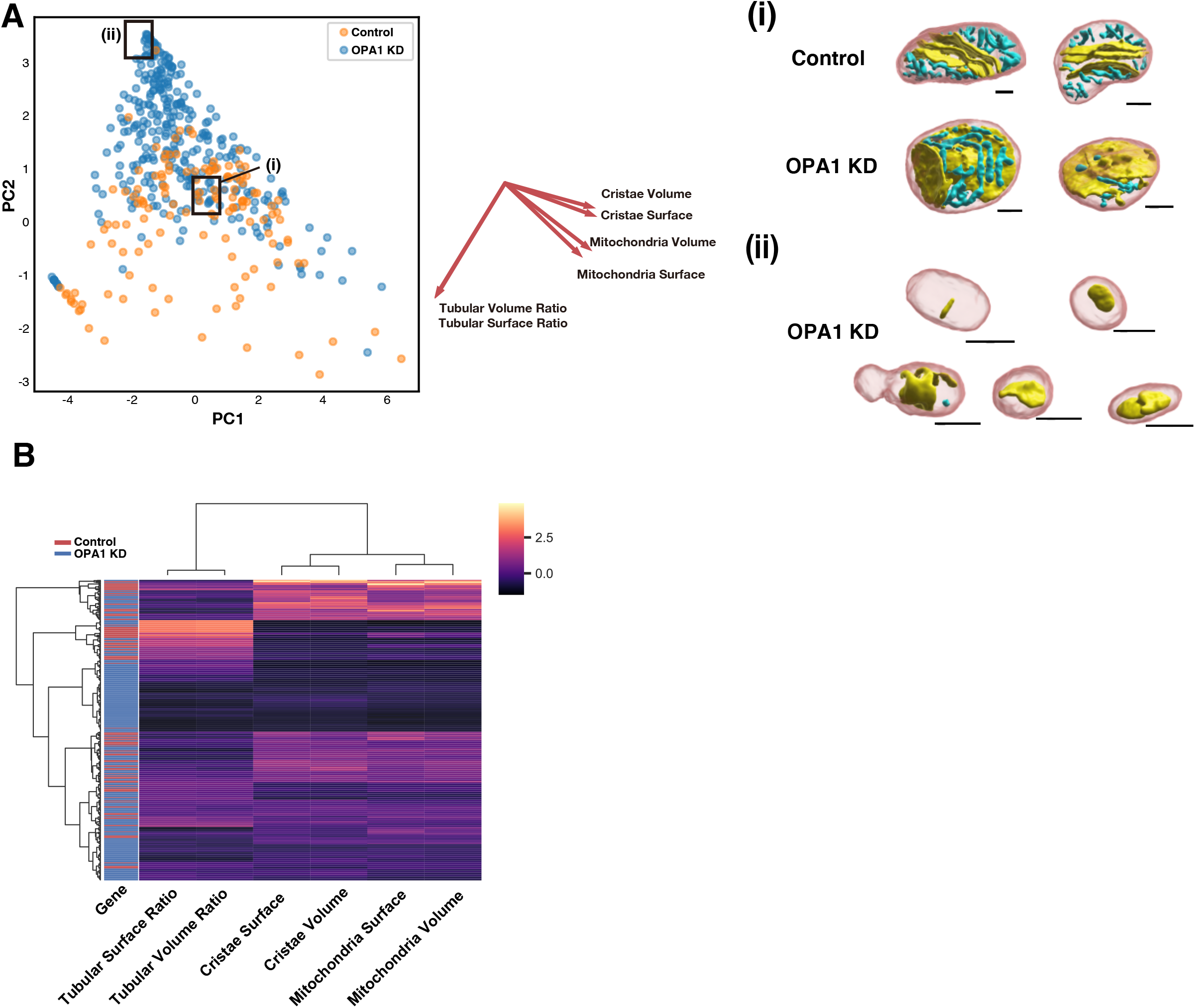
Unsupervised analyses of mitochondria and cristae structure. **(A)** Principal component analysis (PCA) of mitochondria. Parameters of the control mitochondria were used to fit the model. The mitochondria in the control cells are shown as orange dots and mitochondria in OPA1 KD cells are shown as blue dots. Representative reconstructed 3D mitochondrial and cristae structures in the area (i) and (ii) are shown. Scale bar 300 nm. **(B)** Hierarchical clustering using Ward’s method. Clustering revealed that there are a cluster with a high tubular cristae ratio and enriched with control derived mitochondria and a cluster with a low tubular cristae ratio characteristic of OPA1 KD. (Control n = 134; OPA1 KD n = 321)

Next, we performed hierarchical clustering of mitochondria using the Ward’s method (**Figure 5B**). Mitochondria were largely divided by the amount of volume and surface area. Interestingly, within the smaller mitochondria cluster, there was a clearly delineated cluster with high tubular ratio, where the control mitochondria were enriched. In addition, there was a cluster with a very small tubular ratio, most of which were OPA1 KD mitochondria. These results suggest that differences in the ratio of lamellar-tubular cristae best represent the phenotype of OPA1 KD, rather than the mitochondrial size or amount of cristae.

## Discussion

We developed a novel platform for HITL deep learning analysis, PHILOW, and implemented human-in-the-loop-iterative learning as well as the TAP method in the deep learning-based EM image analysis. PHILOW is an open-source python package that provides users with a seamless GUI environment for annotation, data management, model training and prediction, and manual correction.

The accuracy of prediction using HITL-TAP achieved superhuman quality. Indeed, the F1-scores and IoUs between the HITL-TAP prediction and either of the two manual segmentations was larger than that between the two manual segmentations (**Figures 3B and 3C**). The superhuman accuracy of this method was highlighted in the segmentation of tubular cristae structures. Post-hoc analysis showed that human annotators missed around 50% of tubular cristae structures detected by our HITL-TAP method. This demonstrates that the HITL-TAP method implemented by PHILOW is not only fast but also reveals previously neglected complex intracellular structures.

To demonstrate the capability of HITL-learning and TAP method enabled by PHILOW, we performed single mitochondrial cristae analysis at an unprecedented scale for both control and OPA1 KD cells. This analysis allowed us to (1) observe complete cristae structures of a single mitochondrion, (2) perform statistical analysis of mitochondrial structural parameters, (3) reconstruct the thin and tortuous tubular cristae comprehensively, (4) find a rare phenomenon like septa formation in OPA1 KD, which would have been missed with low-throughput analysis, (5) unsupervised clustering of mitochondria purely from the ultrastructure.

Based on observations in control cells, we found multiple important properties of mitochondrial and cristae structures. First, our data indicate that the surface area of cristae per mitochondria volume is 6.6/μm on average. Furthermore, we found that the surface/volume is similar between tubular and lamellar cristae (0.060/nm and 0.052/nm, respectively). Indeed, changes in the ratio between tubular and lamellar structure in OPA1 KD cells did not affect the total surface area of cristae. This suggests that the morphological variety of cristae is more likely to be important for a function such as controlling diffusion of ion species rather than increasing the surface area (Mannella et al., 2013).

3D analysis of large numbers of mitochondria provided us with highly reliable quantitative information about the dimension of mitochondrial structures. For example, our analysis showed that the surface area per mitochondria volume was about 0.006/nm. Although this value is smaller than the values in other cell types (0.02-0.04/nm, Else and Hulbert, 1985; Safiulina et al., 2006; Schwerzmann et al., 1989), a recent study using FIB-SEM reports that it was 0.007/nm in the liver (Murphy et al., 2010). This suggests that the evaluation of morphology highly depends on imaging techniques and sample treatments. Since the chemical fixation adds artifacts to the cellular ultrastructure, further study using cells fixed with high-pressure freezing is required for a more precise evaluation of cristae structures.

Using the HITL-TAP method, we demonstrated that OPA1 is required for the proper formation of CJ and tubular cristae without affecting the total surface area of cristae. OPA1 has been known as a molecular determinant of IMM structures via its GTPase-dependent IMM fusion function and -independent CJ forming functions (Patten et al., 2014). In terms of regulating cristae, our quantitative analysis and 3D inspections strongly support that OPA1 is responsible for CJ formation but only moderately for IMM fusion. In line with this, the onion-like structure is observed in the cells deficient for Mic60 or Mic19, which are required for CJ formation (Stephan et al., 2020). A mathematical modeling suggested that the morphology of cristae determines the diffusion patterns of ions and metabolites (Mannella et al., 2013). More precise simulation using actual 3D structures obtained in this study will reveal the contribution of cristae morphology to physiological events, such as apoptosis (MacVicar and Langer, 2016).

Our workflow allows previously unattainable quantitative analysis of tubular cristae structures in genetically modified cells. Although deep learning algorithms, isotropic FIB-SEM imaging, and high-performance GPUs have been developed individually, it is only the efficient combination of these in our human intervention interface by PHILOW that allows us to observe structures that humans cannot readily recognize. Thus, PHILOW paves the way for unbiased large-scale analyses of the cellular ultrastructure with vastly improved accuracy and efficiency compared to previous approaches. Additionally, PHILOW allows for identification of cellular nanostructures that are not yet recognizable by humans. This pioneering technology will advance not only the field of cell biology, but also allow for capturing subtle phenotypical symptoms occurring in the early stages of diseases at the ultrastructural level of vital organelles.

## STAR★Methods

### KEY RESOURCES TABLE

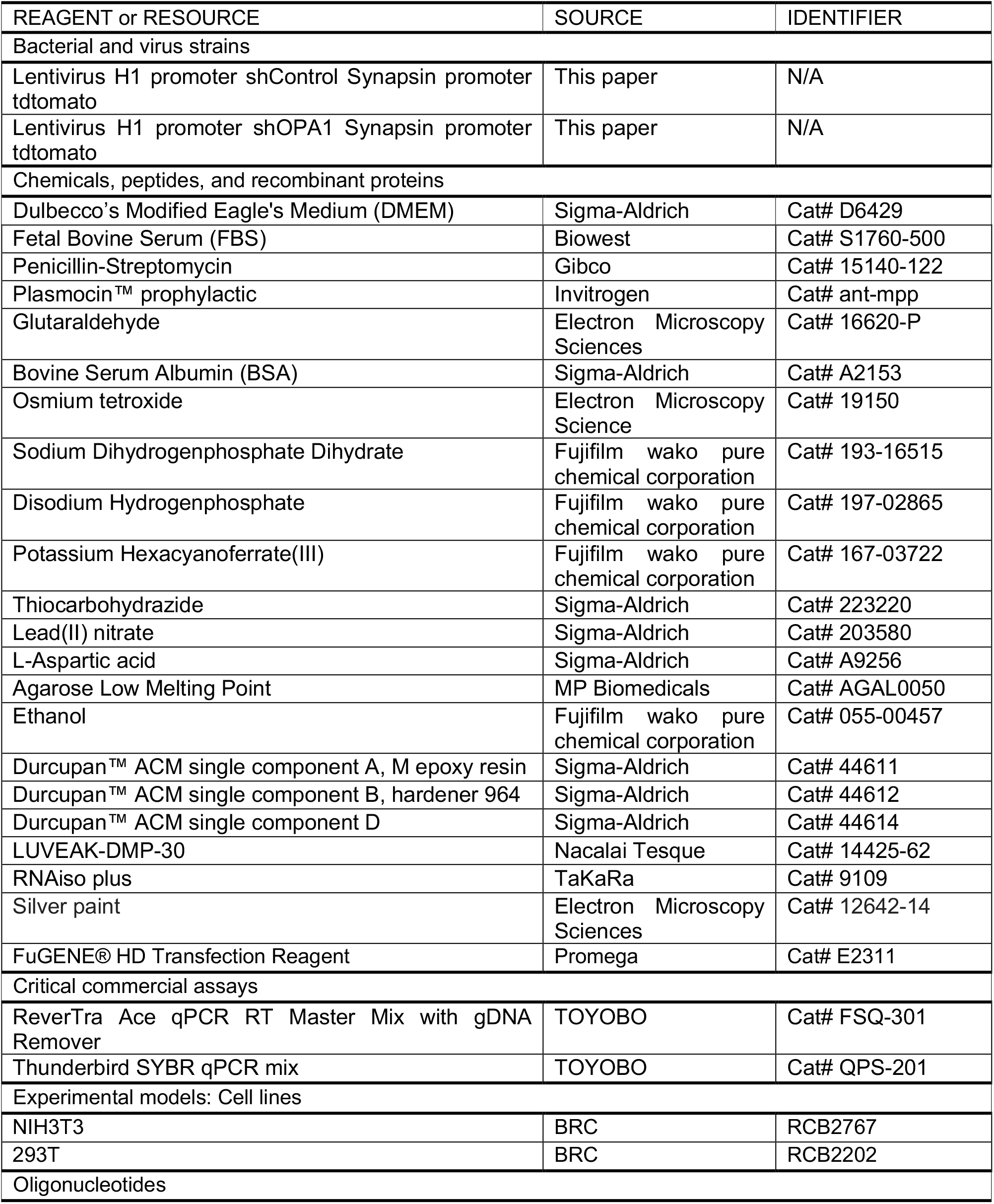
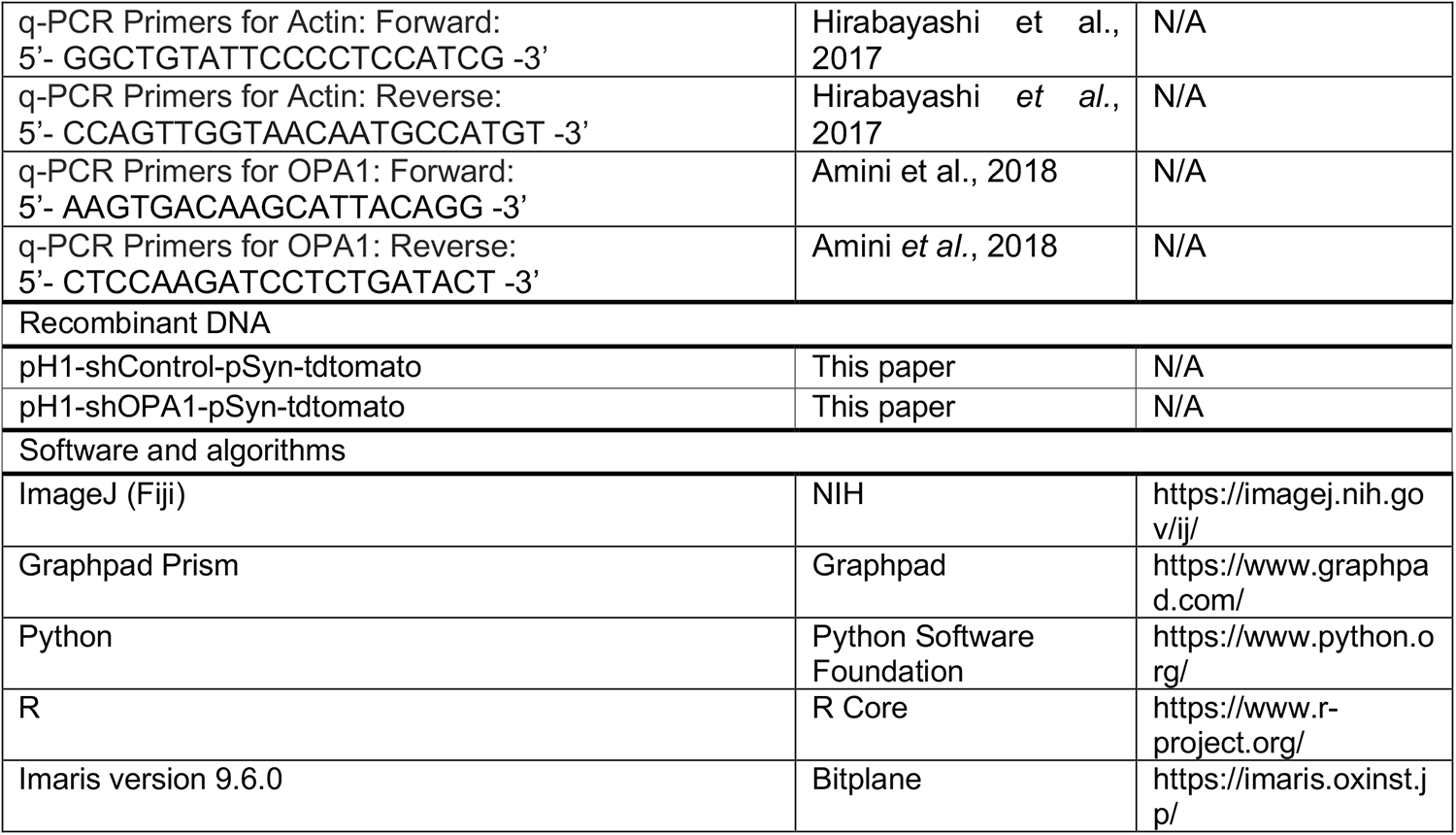

***napari contributors (2019). napari: a multi-dimensional image viewer for python. doi:10.5281/zenodo.3555620***

### RESOURCE AVAILABILITY

#### Lead Contact

Further information and requests for resources and reagents should be directed to and will be fulfilled by the Lead Contact, Yusuke Hirabayashi.

#### Materials Availability

Plasmids generated in this study are being deposited to Addgene.

#### Data and Code Availability

All original code has been deposited at https://github.com/neurobiology-ut/PHILOW/ and is publicly available as of the date of publication.

### EXPERIMENTAL MODEL AND SUBJECT DETAILS

#### Cell culture

NIH3T3 cells were maintained with Dulbecco’s Modified Eagle Medium supplemented with 10% FBS, 1% Penicillin-Streptomycin, and 5 μg/mL Plasmocin™ prophylactic at 37°C under 5% CO_2_.

#### DNA plasmids

pH1-shControl-pSyn-tdtomato and pH1-shOPA1-pSyn-tdtomato were generated from pH1-shSRGAP2-pSyn-tdtomato (kind gift from Dr. Franck Polleux, generated from pSCV2; Charrier et al., 2012) by replacing the shSRGAP2 sequence with hybridized oligomers of shControl (5’-TCG AGC CGC AGG TAT GCA CGC GTT CAA GAG ACG CGT GCA TAC CTG CGG TTT TTG T -3’) and shOPA1 (5’-CCG GCA TGG AAG AAG AAC CAT ATT TCT CGA GAA ATA TGG TTC TTC TTC CAT GTT TTT G -3’), respectively, using XhoI and XbaI sites.

### METHOD DETAILS

#### FIB-SEM

NIH3T3 cells infected with lentivirus carrying either the control or shOPA1 were fixed with 2.5% glutaraldehyde in DMEM, for 1 hour at room temperature. After washing with 0.1 M phosphate buffer (0.02 M Sodium Dihydrogenphosphate Dihydrate, 0.08 M Disodium Hydrogenphosphate) cells were scraped and collected with 0.2% BSA/0.1 M phosphate buffer followed by centrifugation at 1,450×g. The samples were post-fixed with 1% OsO_4_, 1.5% potassium ferricyanide in a 0.1 M phosphate buffer for 2 hour (for samples in Figure 2 and 3) or 30 minutes (for samples in Figure 4 and Figure 5). After being rinsed for 3 times with H_2_O, cells were stained with 1% thiocarbohydrazide for 5 minutes. After rinsing with H_2_O for three times, cells were stained with 1% OsO_4_ in H_2_O for 30 minutes. After rinsing with H_2_O for two times at room temperature and three times with H_2_O at 50°C, cells were treated with 0.635% lead nitrate, 0.4% aspartic acid, pH 5.2 at 50°C for 20 minutes. The final cell pellet was embedded into 2% low melting agarose (MP Biomedicals). After gel setting on ice, the embedded cell pellets were cut into small fragments (1-2 mm^3^). The fragments were followed by incubations in an ascending ethanol series (10 minutes each in 50%, 70%, 90%, 95% ethanol/H_2_O), 10 minutes in 100% ethanol 4 times. This was followed by infiltration in graded concentrations of Durcupan resin (Durcupan-ethanol for 1 day at a 1:3 dilution, 1 day at a 1:1 dilution, and 1 day at a 3:1 dilution). After incubating with 100% Durcupan resin overnight, curing of the resin was achieved at 65°C for 3 days. Durcupan resin were made by mixing 12.4 g of component A, 9.9 g of component B, 0.2 g of LUVEAK-DMP-30, and 0.1 ml of component D. Resin blocks were trimmed with a TrimTool diamond knife (Trim 45; DiATOME) in a Leica Ultramicrotome (UC7). The resin blocks were clued with silver paint (12642-14, Electron Microscopy Sciences) on a SEM stub and coated with gold or platinum to guarantee electrical conductivity. The stubs were fixed on a 45° multi holder (EMS) and mounted in a FIB-SEM (Helios 650, Thermo Fisher). Milling was done at 52°, parallel to the gallium ion beam (30 kV, 770 pA), removing 10 nm material per step. Imaging was done normal to the electron beam, as described previously (Kizilyaprak et al., 2015) at 1.5 keV – 2 keV, 800 pA beam current, 6,144×4,096 frame size, 2.7 mm – 2.8 mm working distance, 30 μm horizontal field of view and 6 μs dwell time, using the through-the-lens detector (TLD) in back-scatter mode (BSE). The final voxel size was 4.88×4.88×10 nm^3^.

#### Lentivirus production

Recombinant lentiviruses were produced as previously reported (Kwon et al., 2016). 293T cells were co-transfected with shuttle vectors, LP1, LP2, and VSV-G using FuGENE transfection reagent (Promega). 24 hours after transfection, the media were exchanged with fresh DMEM supplemented with 10% FBS, 1% Penicillin-Streptomycin, and 5 μg/mL Plasmocin™ prophylactic, and 24 hours later, supernatants were harvested, spun at 500×g to remove debris and filtered through a 0.45 μm filter (Sartorius). The filtered supernatant was concentrated to 100 μl using an Amicon^®^ Ultra-15 (molecular weight cut-off 100 kDa) centrifugal filter device (Merck Millipore Ltd.), which was centrifuged at 4,000×g for 60 minutes at 4°C. The concentrated samples (100 μl) were diluted with 150 μl of PBS and stored at −80°C in 50 μl aliquots.

#### Quantitative RT-PCR

Total RNA was isolated from NIH3T3 cells infected with lentivirus carrying either the control or shOPA1 7 days after infection, with the use of RNAiso plus (TaKaRa), and 0.5 μg of the RNA were subjected to reverse transcription with ReverTra Ace qPCR RT Master Mix with gDNA Remover (TOYOBO). The resulting cDNA was subjected to real-time PCR with Thunderbird SYBR qPCR mix (TOYOBO) in a LightCycler 96 (Roche).

#### Image Pre-processing

Cropped images were aligned using the Linear Stack Alignment with Scale Invariant Feature Transform (SIFT) Plugin, implemented in Fiji. The algorithm was run three times with following parameters: (1) 1.6 pixels Gaussian blur, 3 steps per scale octave, 64 pixels minimum image size, 1,024 pixels maximum image size, 4 feature descriptor size, 8 feature descriptor orientation bins, 0.92 closest/next closet ratio, 25 pixels maximal alignment error, 0.05 inlier ratio, rigid transform and with interpolation; (2) 1.6 pixels Gaussian blur, 3 steps per scale octave, 256 pixels minimum image size, 1,024 pixels maximum image size, 4 feature descriptor size, 8 feature descriptor orientation bins, 0.92 closest/next closet ratio, 25 pixels maximal alignment error, 0.05 inlier ratio, rigid transform and with interpolation; (3) 1.6 pixels Gaussian blur, 3 steps per scale octave, 256 pixels minimum image size, 1,024 pixels maximum image size, 4 feature descriptor size, 8 feature descriptor orientation bins, 0.92 closest/next closet ratio, 25 pixels maximal alignment error, 0.05 inlier ratio, affine transform and with interpolation.

#### Calculation and subsequent analysis of mitochondria and cristae ultrastructural features

First, we labeled mitochondria and counted labeled mitochondrial volume. Then we extracted 1 pixel of each mitochondrial outline and counted them. Next, we calculated the surface area and volume of the cristae in the labeled mitochondria using the same method as for mitochondria. Finally, we calculated the surface area and volume of lamellar and tubular structures by separating whether the calculated cristae were lamellar or tubular. To calculate the angle between mitochondria and lamellar cristae, we calculated the first principal component (PC) of their voxel locations and set its axis as the direction of the mitochondrial longitudinal axis (mitochondrial orientation). For lamellar cristae, lamellar cristae volumes are eroded at first to separate which are in contact with each other. Then we calculated the third PC and set it as the axis perpendicular to the lamellar cristae plane. The angle between the two axes was then calculated. For unsupervised analysis of 3D mitochondria and cristae structures, mitochondria volume, mitochondria surface, cristae volume, cristae surface, tubular cristae volume and lamellar cristae volume were counted as above. Tubular cristae volume and surface ratio were calculated as divided tubular cristae volume by sum of tubular and lamellar cristae volume to calculate tubular cristae ratio. Features were standardized by removing the mean and scaling to unit variance before use in principal component analysis (PCA) and hierarchical clustering using Ward’s method with Euclidean distance. One mitochondrion with deviated size and three mitochondria without cristae were excluded before analysis. More details with the source code and documentation are available at: https://github.com/neurobiology-ut/PHILOW_Data_Manuscript

#### Implementation

The segmentation algorithms were trained using tensorflow 1.13.0, and the quantification algorithms were written in Python 3.7.0 using the libraries: numpy 1.17.4, opencv-python 4.0.0.21, scikit-image 0.17.2, scipy 1.3.2, and napari 0.4.0 (napari contributors, 2019). These algorithms were accelerated using the GPUs (NVIDIA TITAN RTX and NVIDIA A100) on Windows 10 with 3.6 GHz Intel Core i9-9900K processor and 32 GB memory. More details with the source code and documentation are available at: https://github.com/neurobiology-ut/PHILOW

#### 3D Visualization

3D visualization of mitochondrial structure and cristae structure was performed using Imaris version 9.6.0 (Bitplane).

### QUANTIFICATION AND STATISTICAL ANALYSIS

Data were statistically analyzed by Student’s t-test or Mann - Whitney test using Graphpad Prism, and graphs were generated using Graphpad Prism. All other statistics and generating graphs were performed using R, including the ggplot2 package or using Python, including matplotlib 3.3.3 and seaborn 0.11.1 packages.

## Author contributions

H.K., and Y.H. conceptualized the project. S.S., K.N., H.K., and Y.H. designed the experiments. K.N. prepared samples for FIB-SEM. B.M.H. acquired FIB-SEM data. S.S. and H.K. developed machine learning algorithms and performed network training and automatic evaluations. S.S. and H.K. wrote the whole code of PHILOW. S.S. and K.N. provided manual annotations, evaluations, and proofreading. S.S. and H.K. performed quantitative analysis from segmented data. S.S., K.N., B.M.H., H.K., and Y.H. wrote the manuscript. H.K. and Y.H. supervised the project.

## Declaration of Interests

H.K. is an employee of LPIXEL Inc.

## Acknowledgements

We thank Drs. Franck Polleux, Heike Blockus, Tommy L. Lewis, Jr, and Yukiko Gotoh for their critical reading of the manuscript and members of the Hirabayashi lab for constructive discussions.

We would like to thank the members of LPIXEL Inc. for helpful discussions on analysis methods and advice on implementation.

This work was supported by JSPS KAKENHI under Grant Number 20H04898, AMED under Grant number JP21dm0207082, and SECOM Science and Technology Foundation Research grant (Y.H.). BMH acknowledges the generous support of the Okinawa Institute of Science and Technology, the Japanese Cabinet Office and Basis for Supporting Innovative Drug Discovery and Life Science Research (BINDS) from AMED under grant numbers 19am0101116j0003 and 20am0101116j0004.

## Supplementary Figure legends

**Figure S1.**
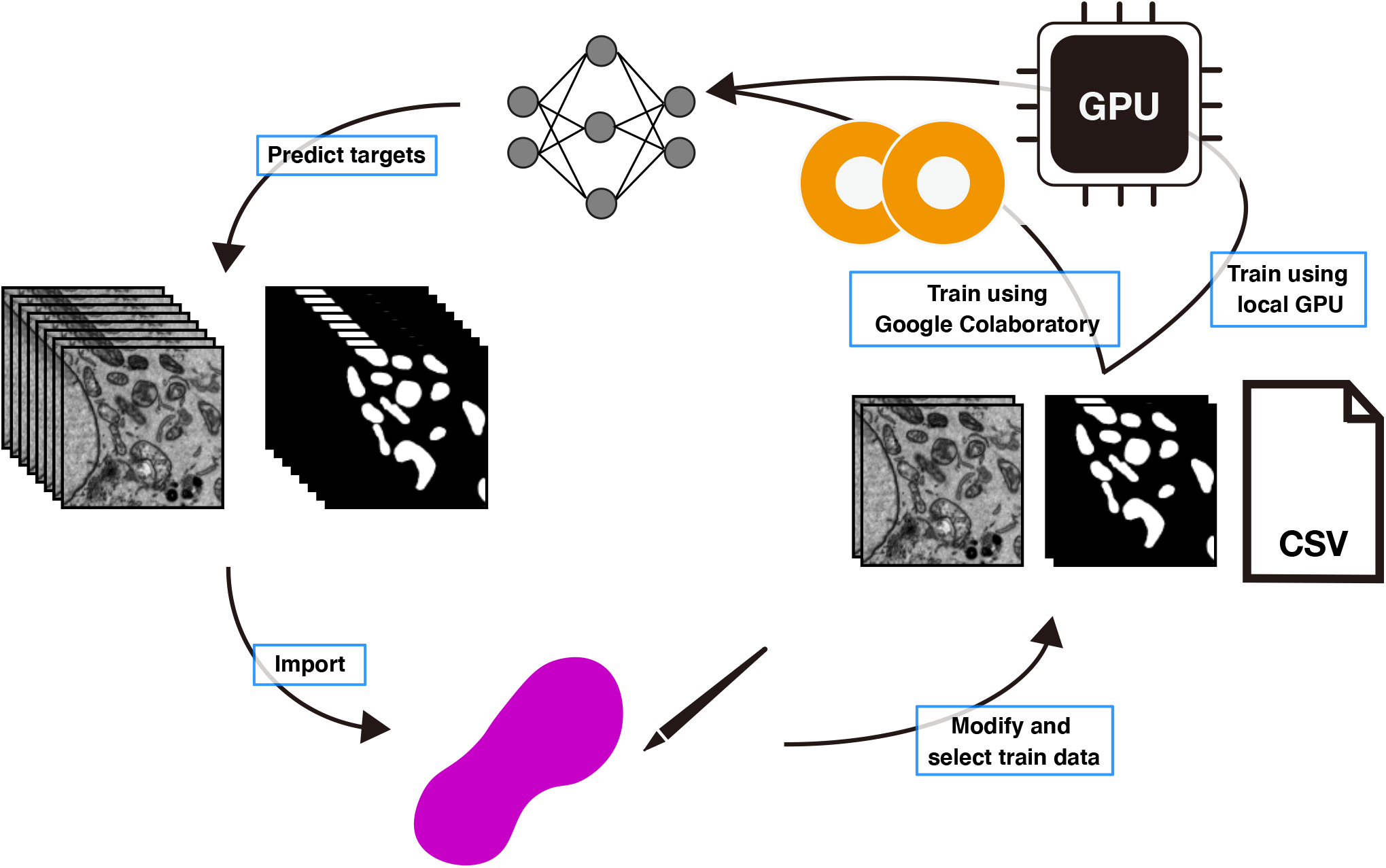
Workflow of the HITL-iterative learning on PHILOW, Related to Figure 1. PHILOW facilitates the HITL-iteration cycle by implementing a segmentation labeling interface, training data management in CSV format, connection to either local GPUs or services in the cloud (Google Colaboratory), and seamless transfer of prediction results back into the segmentation labeling interface.

**Figure S2.**
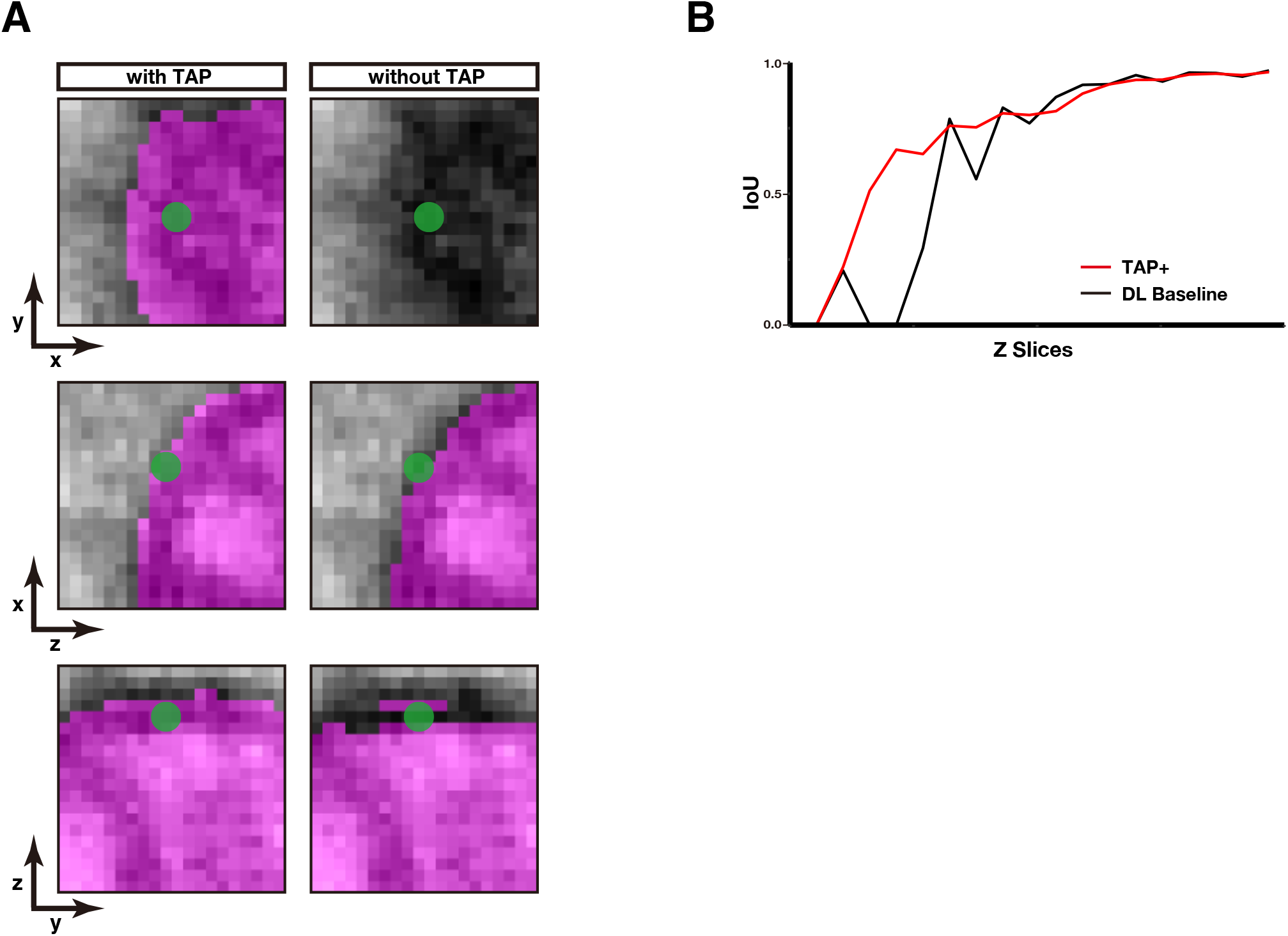
Mitochondrial segmentations at the edges with and without TAP, Related to Figure 2. (A) Segmentations of a mitochondrion with or without the TAP method. The edge of mitochondrion was not annotated as part of mitochondria by a 2D UNet++ based prediction only from the xy plane (without TAP). With the TAP method, the same edges of mitochondrion were successfully annotated as a part of mitochondria by combining the predictions from all three axes. (B) The IoU overlap with the next slice was measured for mitochondria segmentation of each slice in the prediction either using the TAP + DL baseline (red) or only DL baseline (black).

**Figure S3.**
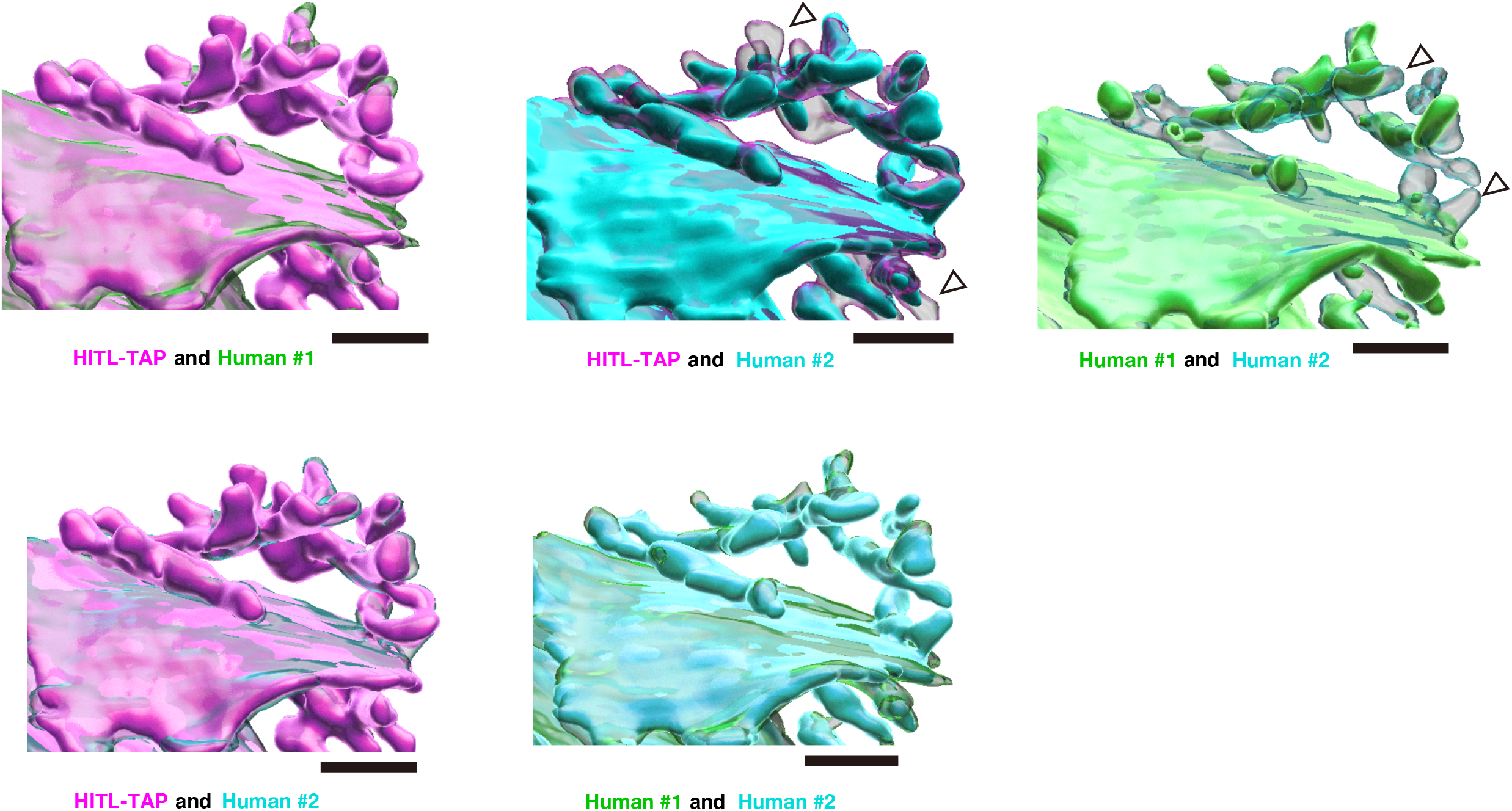
Visualization of the segmentation differences between the HITL-TAP and human annotators, Related to Figure 3. Two of the 3D reconstructions by the HITL-TAP, Human #1 or Human #2 are compared. Note that the human annotators tend to segment lamellar structures more extensively than the HITL-TAP. Also note that the human annotators tend to omit tubular structures, which were recognized by the HITL-TAP (arrowheads). (A) Magenta (filled): HITL-TAP; Green (transparent): Human #1 (B) Cyan (filled): Human #2; Magenta (transparent): HITL-TAP (C) Green (filled): Human #1; Cyan (transparent): Human #2 (D) Magenta (filled): HITL-TAP; Cyan (transparent): Human #2 (E) Cyan (filled): Human #2; Green (transparent): Human #1

**Figure S4.**
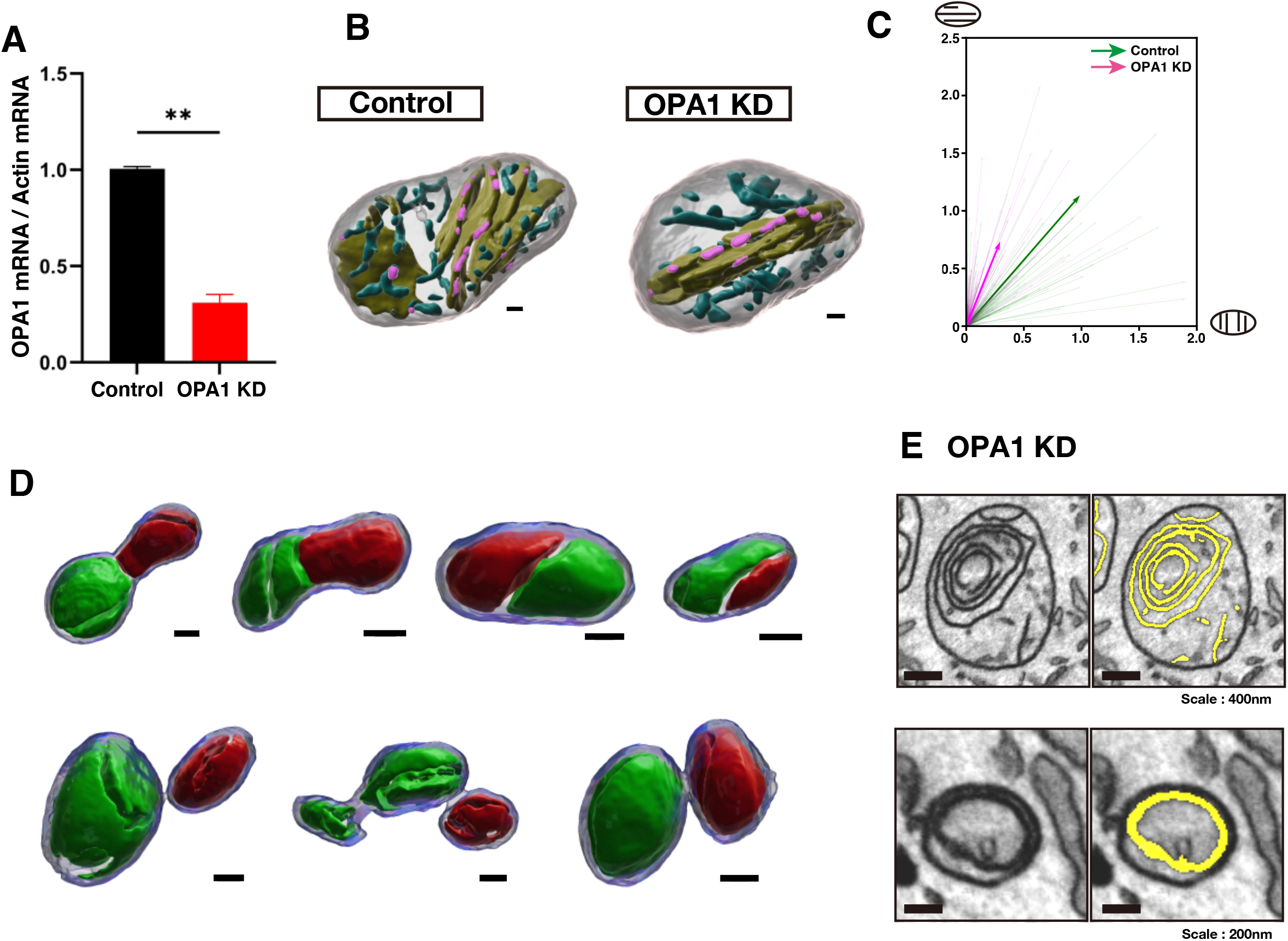
Characteristic morphologies of cristae in OPA1 KD cells, Related to Figure 4. (A) The amounts of OPA1 mRNA were quantified by quantitative RT-PCR and normalized by Actin mRNA levels. Error bars indicate standard errors from duplicated samples of quantitative PCR. **p<0.01, Student’s t-test. (B) Representative mitochondrial structures either from the control or OPA1 KD cell. Note that the OPA1 KD mitochondrion has less cristae junction area than the control mitochondrion. Yellow: lamellar cristae structure; Cyan: tubular cristae structure, Magenta: cristae junctions. Scale bars, 100 nm. (C) The light-colored vectors indicate the angular direction between the vector perpendicular to each lamellar cristae and the mitochondrial long-axis vector. The magnitude was normalized by the inverse of the eigenvalue. The dark colored vectors are the average of vectors from all mitochondria scaled by a factor of 4. Green: Control; Magenta: OPA1 KD. (D) Four OPA1 KD mitochondria with complete septa. Each mitochondrion was separated into the green and red compartments by inner membranes. These compartmentalized mitochondria were defined by a following method; First, mitochondria were eroded by 4 voxels. Then, when a lamellar cristae structure divides mitochondria into two compartments, and the smaller compartment is more than 15% of the total mitochondrial volume, the structure was defined as potential septum. After a visual inspection, four mitochondria were selected. Three mitochondria omitted by the visual inspection were also shown in the second row. Scale bars, 20 nm. (E) Representative EM images of onion-like structures in the OPA1 KD cells. Yellow: segmentations of lamellar structures.

**Figure S5.**
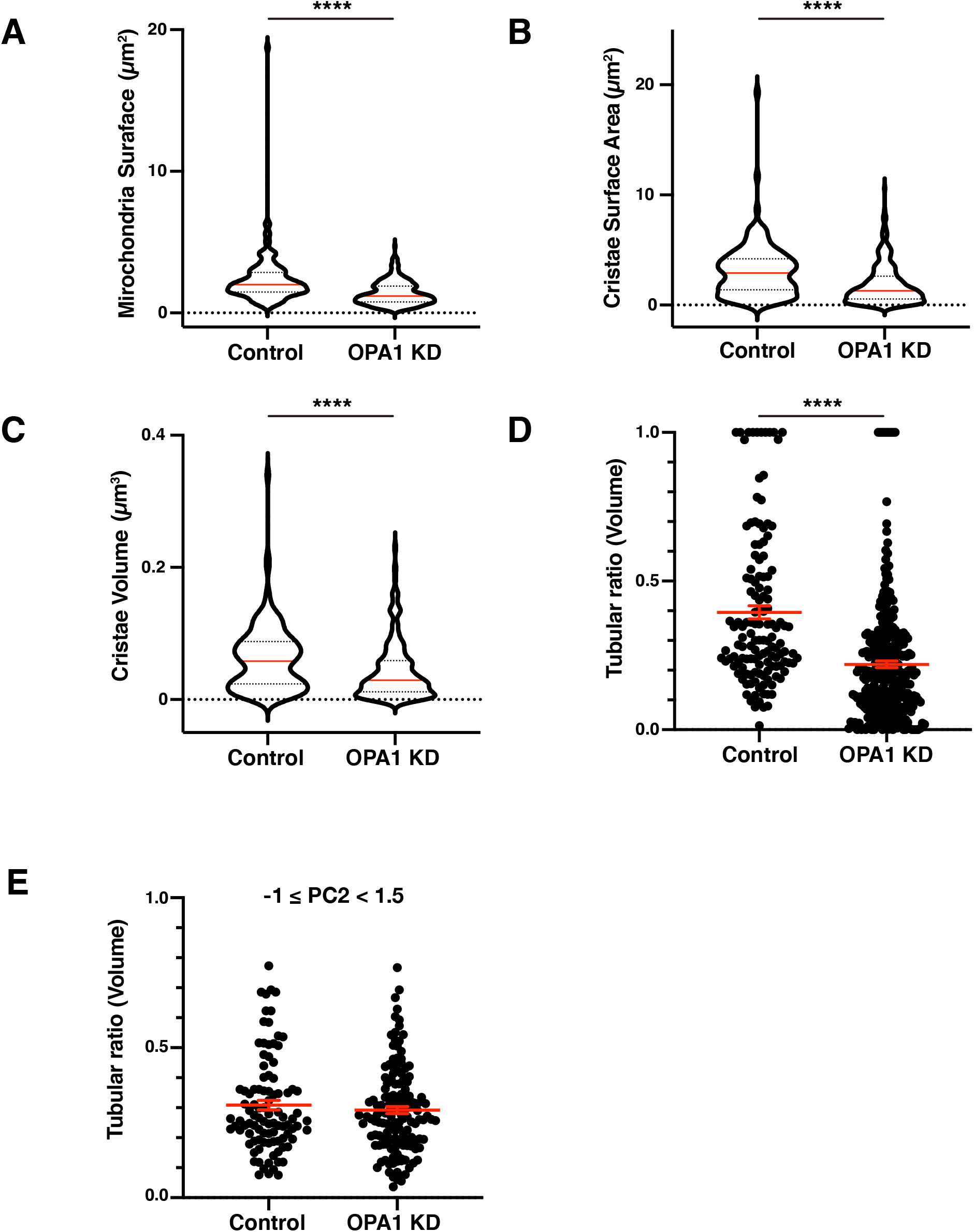
Statistical analyses of each parameter used in the unsupervised analysis, Related to Figure 5. (A-D) Parameters of mitochondria and cristae from the control or OPA1 KD cells are indicated. In addition to these parameters, mitochondrial volume (Figure 4C), and Tubular ratio (surface, Figure 4E) were used for the PCA and hierarchical analysis. Red lines show median in (A)-(C) and mean ± SEM in (D). (E) Tubular volume ratio between the control and the OPA1 KD mitochondria within the range of −1 ≤ PC2 < 1.5. There was no significant difference between the control and OPA1 KD. ****p < 0.0001, Mann–Whitney test.

## Supplementary Video legends

**Video S1. Representative 3D ultrastructure of the control mitochondria and cristae reconstructed using HITL-TAP on PHILOW, related to Figure 2.**

Mitochondria and cristae were successfully segmented from the FIB-SEM stack using HITL-TAP on PHILOW. Cristae were also categorized into lamellar structure and tubular structure by HITL-TAP on PHILOW. Magenta or white: mitochondrial outer membrane, yellow: lamellar structure, cyan: tubular structure.

**Video S2. Segmentation differences between the HITL-TAP and the human annotator, related to Figure 3.**

The human annotator missed to segment tubular cristae structures that were well segmented by HITL-TAP method. Green: human annotator, Magenta: HITL-TAP.

**Video S3. Representative 3D ultrastructure of the OPA1 KD mitochondria and cristae reconstructed using HITL-TAP on PHILOW, related to Figure 4.**

OPA1 KD mitochondria and cristae were successfully segmented from the FIB-SEM stack using HITL-TAP on PHILOW. Cristae were also categorized into lamellar structure and tubular structure by HITL-TAP on PHILOW. Magenta or white: mitochondrial outer membrane, yellow: lamellar structure, cyan: tubular structure.

**Video S4. Comparison of representative 3D ultrastructure of mitochondria between the control and OPA1 KD cells, related to Figure 4.**

3D ultrastructure of three mitochondria each from the control cell and the OPA1 KD cells. Note that the tubular structure was significantly less in OPA1 KD mitochondria than the control ones. White: mitochondrial outer membrane, yellow: lamellar structure, cyan: tubular structure.

**Video S5. Representative 3D ultrastructure of mitochondria containing onion-like structures in OPA1 KD cells, related to Figure 4.**

Representative 3D reconstruction of an onion-like mitochondrion. Note that the onion-like structure does not form a septum. Some parts of the inner membrane are fused to the outer membrane. White: mitochondrial outer membrane, yellow: lamellar structure, cyan: tubular structure.

## Notes

https://github.com/neurobiology-ut/PHILOW

https://github.com/neurobiology-ut/PHILOW_Data_Manuscript

